# Differentiation of primate primordial germ cell-like cells following transplantation into the adult gonadal niche

**DOI:** 10.1101/226506

**Authors:** Enrique Sosa, Di Chen, Ernesto J. Rojas, Jon D. Hennebold, Karen A. Peters, Zhuang Wu, Truong N. Lam, Jennifer M. Mitchell, Ramesh C. Tailor, Marvin L. Meistrich, Kyle E. Orwig, Gunapala Shetty, Amander T. Clark

## Abstract

A major challenge in stem cell differentiation validation is the availability of bioassays to prove cell types generated *in vitro* are equivalent to cells *in vivo*. In the mouse model, differentiation of primordial germ cell-like cells (PGCLCs) from pluripotent cells was validated by transplantation, leading to the generation of spermatogenesis and to the birth of offspring. Here we report the use of xenotransplantation (monkey to mouse) and homologous transplantation (monkey to monkey) to validate our *in vitro* protocol for differentiating male rhesus macaque PGCLCs (rPGCLCs) from rhesus macaque induced pluripotent stem cells (riPSCs). Specifically, transplantation of aggregates containing rPGCLCs into mouse and nonhuman primate testicles overcomes a major bottleneck in rPGCLC differentiation with the expression of VASA and MAGEA4, but not ENO2. These findings suggest that immature rPGCLCs once transplanted into an adult gonadal niche commit to differentiate towards late PGCs and spermatogonia-like cells but do not complete the conversion into ENO2-positive spermatogonia.

## Introduction

Germline cells are essential for fertility and passing DNA from one generation to the next. In each generation, germ cell development begins around the time of embryo implantation with the differentiation of founding progenitors called primordial germ cells (PGCs). PGCs are transient and in the appropriate environment will subsequently advance in differentiation towards oogonia in females and pro-spermatogonia in males.

In an inappropriate environment, however, the latent pluripotency program can be reactivated leading to germ cell tumors including teratomas. Moreover, abnormal differentiation of PGCs has the potential to impact the quality of the entire cohort of germ cells in the adult gonad given that after PGC specification no other cell type can contribute to the germline. Therefore, understanding the biology of PGCs has important implications for future reproductive success and child health.

One of the most exciting models for understanding human PGC development is the pluripotent stem cell model and differentiation into PGC-like cells (PGCLCs) *in vitro*^1,2,3,4^. Directed differentiation protocols for generating human PGCLCs (hPGCLCs) result in the formation of so called “early PGCs” which are equivalent to PGCs at around week 3 of human embryo development. Early PGCs in the primate cynomolgus (cyno) macaque are triple positive for SOX17, PRDM1 and TFAP2C, while being negative for the “late stage” PGC markers VASA and DAZL^5^. A recent study has demonstrated that female human embryonic stem cell (hESCs) can differentiate into VASA-positive human oocyte-like cells^6^. However, an approach for differentiating male primate PGCLCs into more advanced VASA positive stages is lacking.

Advanced differentiation and generation of fertilization competent sperm from mouse PGCLCs (mPGCLCs) was first shown by transplantation of mouse aggregates and mPGCLCs into the testicles of infertile male mice^7, 8, 9^. Furthermore, mPGCLCs have been differentiated entirely *in vitro* using co-culture with gonadal somatic cells^10^. The differentiation of male mPGCLCs entirely *in vitro* depended first upon the success of testicular transplantation to prove mPGCLC competency. In humans transplanting hPGCLCs into the testicles of human subjects as a first-line experiment to prove hPGCLC competency is inconceivable. Instead, we propose that a first approach could instead utilize the testicular xenotransplantation bioassay, or alternatively homologous transplantation of nonhuman primate PGCLCs.

Testicular xenotransplantation involves transplantation of primate (human or nonhuman) testicular cells containing germ cells into the seminiferous tubules of busulfan-treated or irradiated immune deficient nude mice^11,12,13,14,15^. More recently, it was also shown that rhesus macaque PGCs (rPGCs) and human PGCs (hPGCs) can also persist and form colonies at the basement membrane of this model, indicating that the testicular xenotransplantation approach can be extended to characterize less mature germline cells, and possibly PGCLCs^16^. In all reported cases of xenotransplantation, human and nonhuman primate germ cells do not differentiate into haploid sperm in the mouse seminiferous tubule niche. Instead, they recapitulate many of the characteristics that are unique to male germline stem cells. These include the ability to 1) migrate to the basement membrane of seminiferous tubules, 2) divide to produce chains of cells with spermatogonial characteristics (a high nuclear to cytoplasmic ratio and intercellular bridges) and 3) persist for long periods of time.

In order to confirm that the testicular xenotransplantation bioassay could be used as an important reporter for germline competency despite the lack of apparent differentiation, Hermann and colleagues^17,18^ subsequently reported that homologous transplantation into recipients depleted of spermatogonial stem cells prior to transplantation can result in complete spermatogenesis from donor cells^15,19^. Furthermore, not only were the donor SSCs competent to undergo complete spermatogenesis the donor-derived sperm were competent to fertilize rhesus macaque oocytes and give rise to donor-derived embryos^19^. It is unknown how less mature rhesus macaque germ cell types will respond in this assay.

In the current study, we differentiated riPSCs to early (VASA negative) rPGCLCs that we characterize as being similar to bona fide embryonic PGCs in the rhesus macaque that are younger than 28 days of post-fertilization development. Following xenotransplantation into irradiated nude mice or homologous transplantation into irradiated rhesus macaques, we show that the seminiferous tubules environment supports survival of rPGCLCs and promotes further differentiation to a VASA-positive state. Taken together, transplantation to the seminiferous tubule environment promotes rPGCLC differentiation beyond what can currently be achieved *in vitro*.

## Results

### Rhesus macaque PGCLCs correspond to early rPGCs younger than day 28 of embryo development

To date, all directed differentiation strategies to generate primate PGCLCs creates VASA negative cells equivalent to early stage PGCs^1,2^. In the cynomolgus macaque, VASA protein expression is induced in some PGCs at around day 28^5^. To evaluate this in rhesus macaque embryo development we collected time-mated embryonic day 28 (D28) rhesus embryos (n=3) (Fig. 1a). Examination of transverse histological sections in the region of the aorta-gonad-mesonephros (Fig. 1a, box) revealed gonadal ridge epithelium, (Fig. 1b, arrow heads), but no definitive gonad. Immunofluorescence (IF) staining for the transcription factor SRY-box 17 (SOX17) a classical visceral endoderm marker that is also required for hPGC specification^1^ revealed that several subsets of SOX17 positive cells can be found in the dorsal aorta (da), the mesonephros (m), the genital ridge epithelium (Fig. 1c, arrow heads), and the dorsal mesentery (white arrow) (Fig. 1c). IF staining of the transcription factor AP-2 gamma (TFAP2C), a second marker of primate PGCs, revealed that TFAP2C positive cells are restricted only to the dorsal mesentery and the genital ridge epithelium (Fig 1d, box and arrow), and not the dorsal aorta. This is consistent with SOX17 expression in hemogenic endothelium in the dorsal aorta^20^, and TFAP2C being a more discriminating marker of primate PGCs^5^. Triple IF staining of SOX17, TFAP2C, and the transcription factor PR/SET domain 1 (PRDM1), all markers of early PGCs, identifies triple positive rPGCs in the dorsal mesentery, as well as rPGCs closely associated with the gonadal ridge epithelium (white arrow head) (Fig. 1d). Furthermore, all TFAP2C positive rPGCs at D28 co-express the pluripotent transcription factor OCT4 (Fig. 1e). To determine whether rPGCs at D28 correspond to early PGCs (VASA negative) or late PGCs (VASA positive)^5^, we stained for VASA and the rPGC marker TFAP2C. Our data indicate that the majority of D28 PGCs were located in either the dorsal mesentery or the genital ridge epithelium, and the majority of TFAP2C positive rPGCs were also VASA positive (Supplementary Fig. 1a-b). In addition, rare VASA negative TFAP2C positive cells were also identified (Fig. 1e, white arrows). As expected SOX2, is not expressed in any TFAP2C positive rPGCs (Fig. 1e). Once we detected endogenous rPGCs that are transitioning from the early to late stages, we next examined the expression of these transcription factors with differentiation of riPSCs into rPGCLCs. The rPGCLC differentiation strategy involves a modification of the two-step differentiation protocol first described by Sasaki and colleagues^2^. The first step involves harvesting undifferentiated riPSCs cultured on mouse embryonic fibroblasts (MEFs) as single cells, and differentiating the riPSCs for 24 hours to create incipient mesoderm-like cells (iMeLCs) (Fig. 2, Supplementary Fig. 2a). The second step involves differentiating the iMeLCs as three-dimensional aggregates in low adhesion 96-well plates (Fig. 2a). Rhesus PGCLCs are formed in step 2 within the aggregates in response to bone morphogenetic protein 4 (BMP4). In the first experiment, we used IF to test whether the transcription factors SOX17, PRDM1 or TFAP2C are expressed in undifferentiated riPSCs, or iMeLCs prior to aggregate differentiation. We also examined expression of BRACHYURY (BRA), as well as pluripotency transcription factors OCT4 and SOX2, to confirm iMeLC induction. We found that both riPSCs and riMeLCs expressed OCT4 and TFAP2C, and in addition, riMeLCs also expressed BRA as expected^2^. SOX2 was uniformly expressed in riPSCs and heterogeneously in iMeLCs. Importantly, we found that riPSCs, and iMeLCs did not express SOX17 or PRDM1 (Supplementary Fig. 2b-c). We hypothesize these transcription factors could be used together to document emergence of nascent rPGCLCs in the aggregate.

**Fig. 1.**
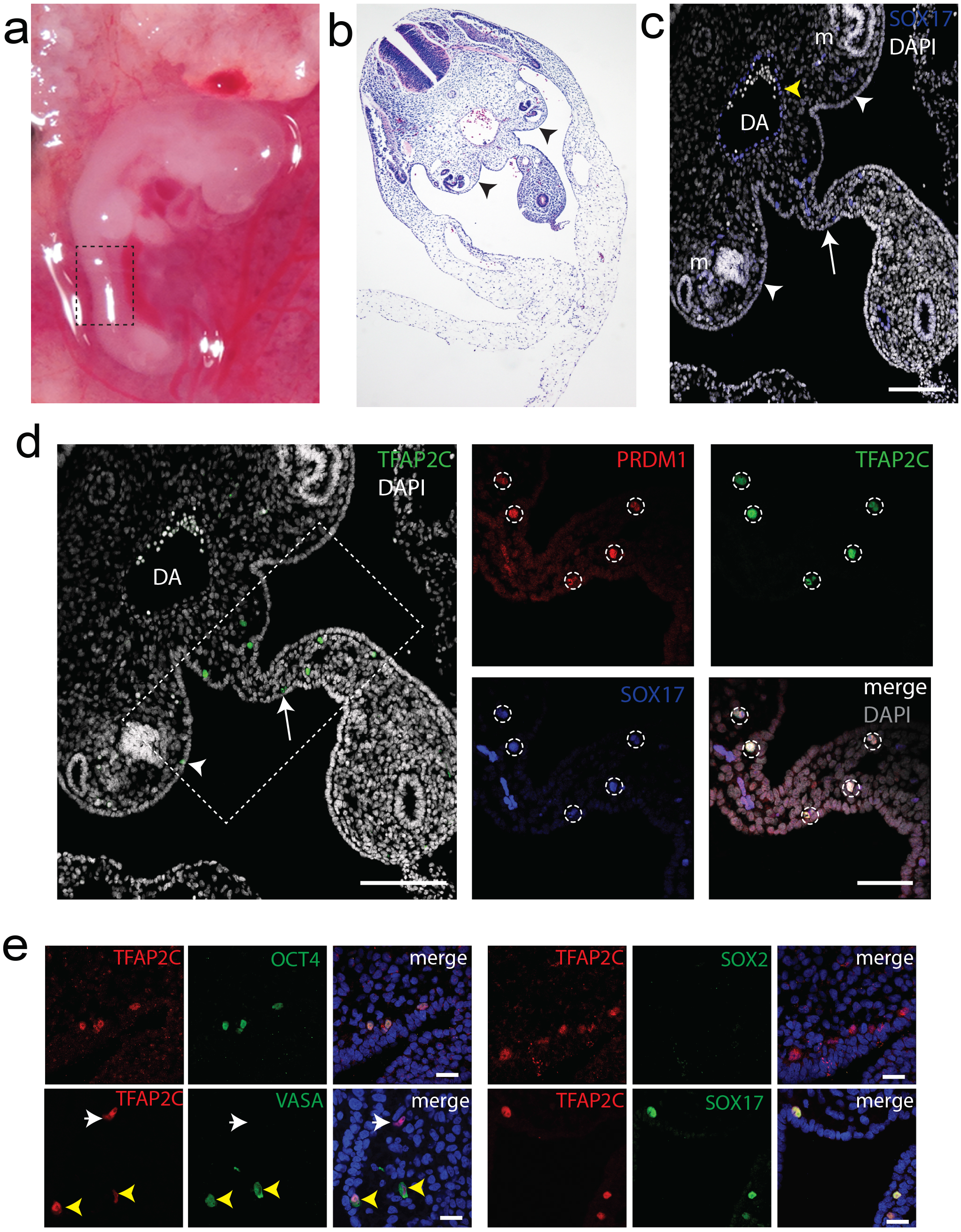
Identification of *in vivo* PGCs in rhesus macaque embyros by immunofluorescence staining of early germ cell markers. (a) Migrating PGCs in a day 28 (D28) rhesus macaque embryo is denoted by the box. (b) Transverse histological section of D28 embryo reveals the genital ridge epithelium (arrow heads). (c) Immunofluorescence staining of D28 sections showing SOX17 (blue) and DAPI nuclear stain (white) in genital ridge epithelium (white arrow heads), dorsal mesentery (white arrow), or hemogenic endothelium (yellow arrow head) within the dorsal aorta (da) along the aorta-gonad-mesonephros of a D28 rhesus macaque embryo. Scale bar, 200 μm. (d) IF staining of TFAP2C discriminates PGCs (box) in a D28 embryo in dorsal mesentery (arrow) as well as the genital ridge epithelium (arrow head). Triple IF staining of PRDM1 (red), SOX17 (blue), and TFAP2C (green) indicates PGCs (dotted circles) express all three markers (merge, white; DAPI nuclear stain, gray). Scale bar, 200 μm. (e) Dual IF of D28 embryos for TFAP2C (red) confirms co-expression with the germ cell marker OCT4, SOX17, and VASA (yellow arrow heads indicate TFAP2C+VASA+ PGCs and white arrow heads indicate TFAP2C+VASA-PGCs), but not SOX2 (green). Merged images show DAPI nuclear stain (blue). Scale bars, 15 μm.

**Fig. 2.**
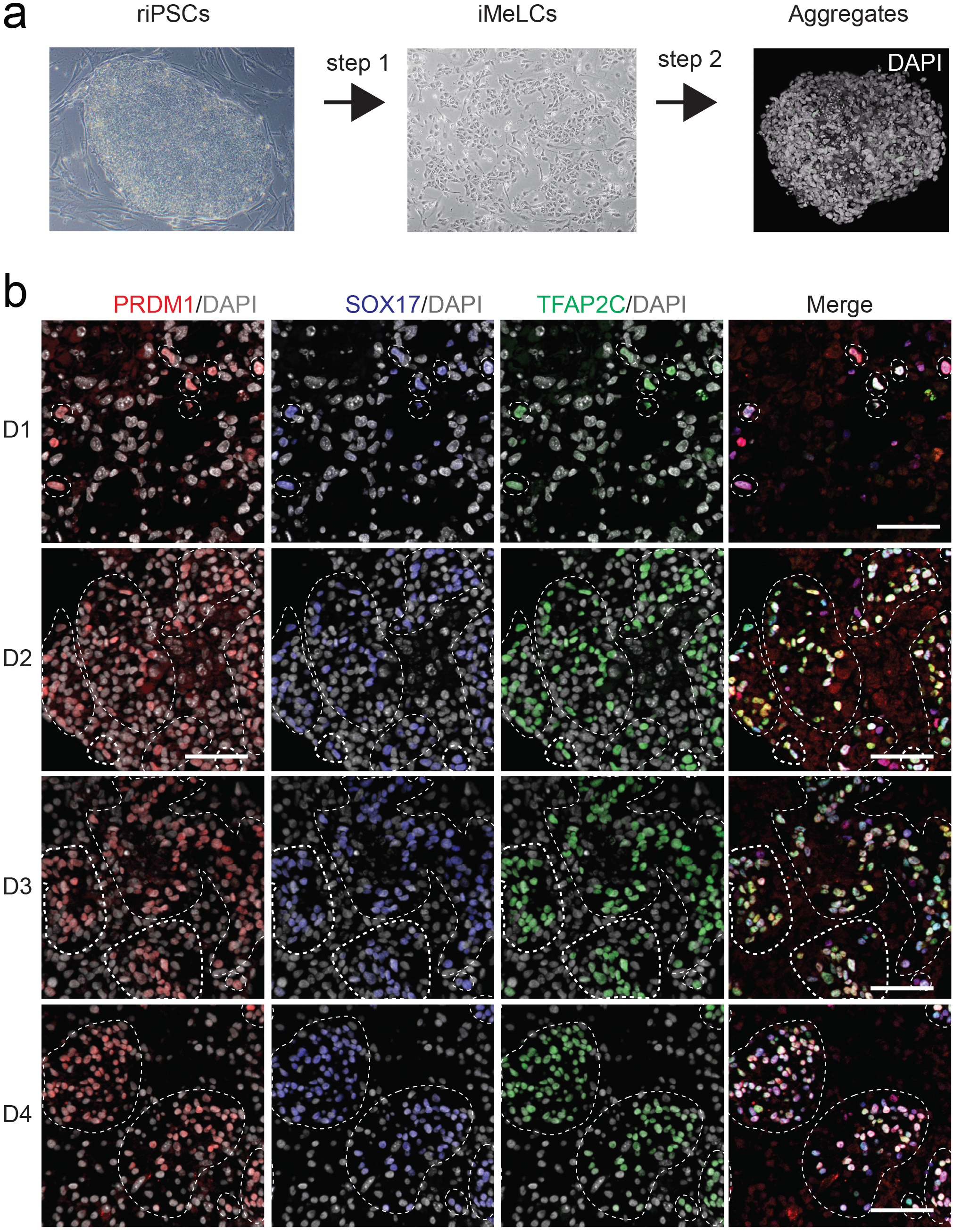
*In vitro* specification of rPGCLCs using pluripotent riPSCs. (a) Schematic overview of rPGCLC-specification in aggregates from riPSCs via an iMeLC intermediate. (b) Triple IF staining for germ cell markers during rPGCLC-specification from Day 1 through Day 4. PRDM1 (red), SOX17 (blue), TFAP2C (green), DAPI (grey) Scale bars, 100μm.

To identify rPGCLCs, aggregates were assessed at day 1 to day 4 for PRDM1 and SOX17 expression. As early as day 1, SOX17 and PRDM1 double positive cells can be identified with only a rare occurrence of single positive cells (Supplementary Fig. 3a). Previous studies analyzing cynomolgus macaque embryos at the time of lineage specification reveals that SOX17/PRDM1 double positive cells mark visceral endoderm (VE) cells, as well as PGCs^5^. Therefore, to discriminate between these two possibilities *in vitro*, we stained SOX17 together with TFAP2C, which is negative in VE and positive in cynoPGCs^5^. Double IF staining revealed that subpopulations of SOX17 positive cells are also positive for TFAP2C at day 1 of differentiation, with more double positive cells identified from day 2 on (Supplementary Fig. 3b).

Based upon the SOX17/TFAP2C dual staining results, we examined rPGCLC formation by analyzing co-expression of PRDM1/SOX17/TFAP2C. Using this approach, we discovered nascent triple positive rPGCLCs as early as day 1 of aggregate differentiation, persisting over the next 8 days. However, by day 15 of aggregate differentiation, rPGCLCs were undetectable (Fig. 2b and Supplementary Fig. 3c-e). In order to confirm that these results were not cell-line dependent, we examined the second iPSC line riPSC90^21^ (Supplementary Fig. 3c-e). Taken together, our results indicate that rPGCLCs are induced at day 1 of aggregate differentiation and can be identified by triple staining for PRDM1/SOX17/TFAP2C.

### Rhesus macaque PGCLCs arrest prior to epigenetic reprogramming

To identify the transcriptional identity of rPGCLCs, we isolated PGCLCs using fluorescence activated cell sorting (FACS) at day 4 and 8 of differentiation using antibodies that recognize EpCAM and ITGA6 (Fig. 3a) and examined the expression of PGC markers. Notably, expression of key early stage PGC markers such as NANOS3, TFAP2C, PRDM1, and cKit, were upregulated in sorted PGCLCs as compared to the parental undifferentiated riPSCs line or riMeLCs (Fig. 3b). To confirm the early stage of rPGCLC development in the aggregates, we examined VASA protein expression in day 4 and day 8 aggregates using IF together with OCT4 as the rPGCLC marker. We discovered that all OCT4 positive rPGCLCs are VASA negative at day 4 and day 8 of differentiation, whereas the positive control D28 rPGCs were mostly VASA positive (Fig. 3c). To evaluate DNA methylation reprogramming we performed IF using antibodies that recognize 5-methyl cytosine (5mC) together in OCT4 positive rPGCLCs. 5mC is detected in early cynoPGCs at the time of specification, but is globally removed from the genome of cynoPGCs at around day 21 of embryo development^5^. Here we show that OCT4 positive PGCs in rhesus embryos at D28 are negative for 5mC as expected. In contrast, the OCT4 positive rPGCLCs at day 4 and 8 of aggregate differentiation were all 5mC positive (Fig 3d). Taken together, the rPGCLCs generated *in vitro* from riPSCs are newly specified, and epigenetically younger than rPGCs found in rhesus macaque embryos at day 28 of embryo development.

**Fig. 3.**
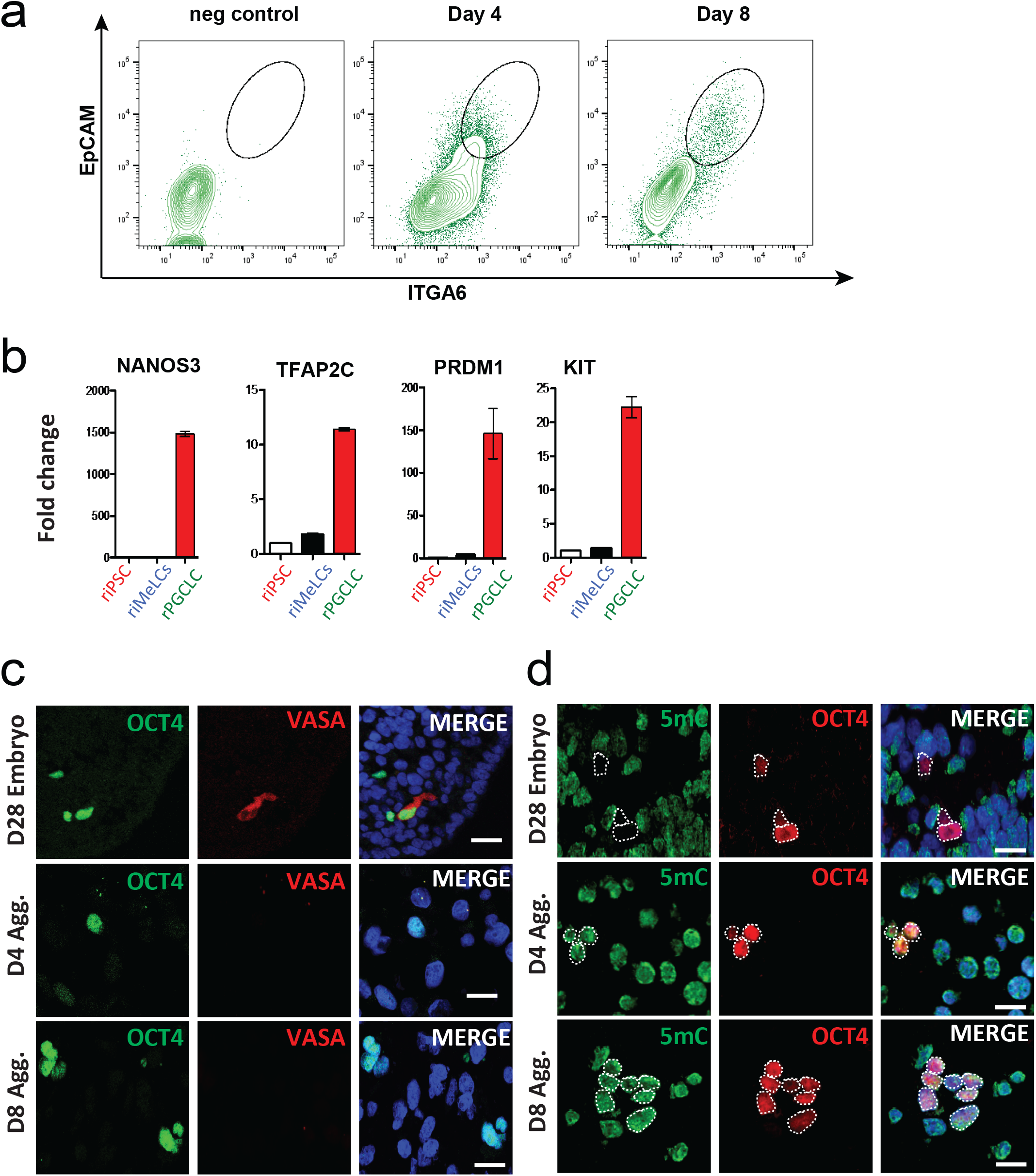
rPGCLCs correspond to early stage PGCs. (a) FACS plots showing sorted EpCAM+ITGA6+ cells in undifferentiated riPSCs from Day 4 (D4) and Day 8 (D8) aggregates. (b) RT-PCR analysis of early stage PGC markers showed transcripts were highly expressed in rPGCLCs, but not riPSCs, or riMeLCs. (c) IF staining of OCT4+ germ cells revealed the late stage PGC marker VASA (red) was only found in D28 embryos (n=3), but not D4 and D8 aggregates. Scale bars, 10μm. (d) IF staining showing expression of the marker of methylation status 5mC was absent from OCT4+ germ cells in D28 embryos (n=3), but present in both D4 and D8 aggregates. Scale bars, 10μm.

### Xeno- and homologous transplantation of early rPGCLCs into the adult gonadal niche promotes differentiation

Previously, it was reported that Carnegie Stage 23 rPGCs xenotransplanted into the seminiferous tubules of the busulfan-treated mouse testis colonize the seminiferous tubule basement membrane giving rise to colony-like chains of cells^16^. To determine whether rPGCLCs persist in this assay and form colonies we first created a green fluorescent protein (GFP) expressing subline of riPSC89 using a lentivirus (referred to as riPSC89^UbiC:GFP^). Prior to transplantation, karyotype analysis was performed to confirm that it possessed a normal 42XY male karyotype consistent with the parent line^22^. Prior to differentiation for xenotransplantation, day 8 aggregates were analyzed by flow cytometry for ITGA6/EpCAM to confirm PGCLC differentiation in the GFP labeled subline. Next, riPSC89^UbiC:GFP^ cells were differentiated through the two-step protocol ending at day 8 of differentiation. Day 8 aggregates were shipped overnight on cold packs to MD Anderson Cancer Center in Houston where they were dissociated and xenotransplanted into the seminiferous tubules of adult immune-deficient nude mice that had been irradiated to ablate endogenous spermatogenesis. The recipient mouse testes were seeded with between 5.0 x 10^4^ to 1.3 x 10^5^, unsorted aggregate cells per transplant (n=6 testicles). As a control, we also transplanted 1.9 x 10^5^-2.7 x 10^5^ undifferentiated riPSCs per transplant (n=6 testicles). Eight weeks after transplant the mice were euthanized, and the testicles were analyzed.

At the time of dissection, 6/6 recipient testes that received a single cell suspension of aggregate cells were GFP positive. In contrast, only 4/6 recipient testes that received undifferentiated riPSCs were GFP-positive (Supplementary Fig. 4a). This result is consistent with previous reports that single cell suspensions of primate pluripotent stem cells exhibit poor survival^23^, which our data would suggest extends to the seminiferous tubule environment. Whole mount IF of the GFP positive testes using anti-rhesus macaque testis cell antibodies^17^ followed by AlexFluor488 secondary antibodies demonstrated that the NHP/GFP positive cells in the recipient testes that received aggregates were VASA positive indicative of rPGCLC differentiation to more mature stages. In contrast, the recipient testes that received undifferentiated riPSCs were VASA negative (Fig. 4a). Furthermore, histological analysis of the GFP-positive recipient testes that received riPSCs yielded teratomas in 2 out 4 of cases. Recipient testes that received rPGCLCs did not form teratomas. Cysts were noted, however, in 4/6 testes (Supplementary Fig. 4b-e, asterisk).

**Fig. 4.**
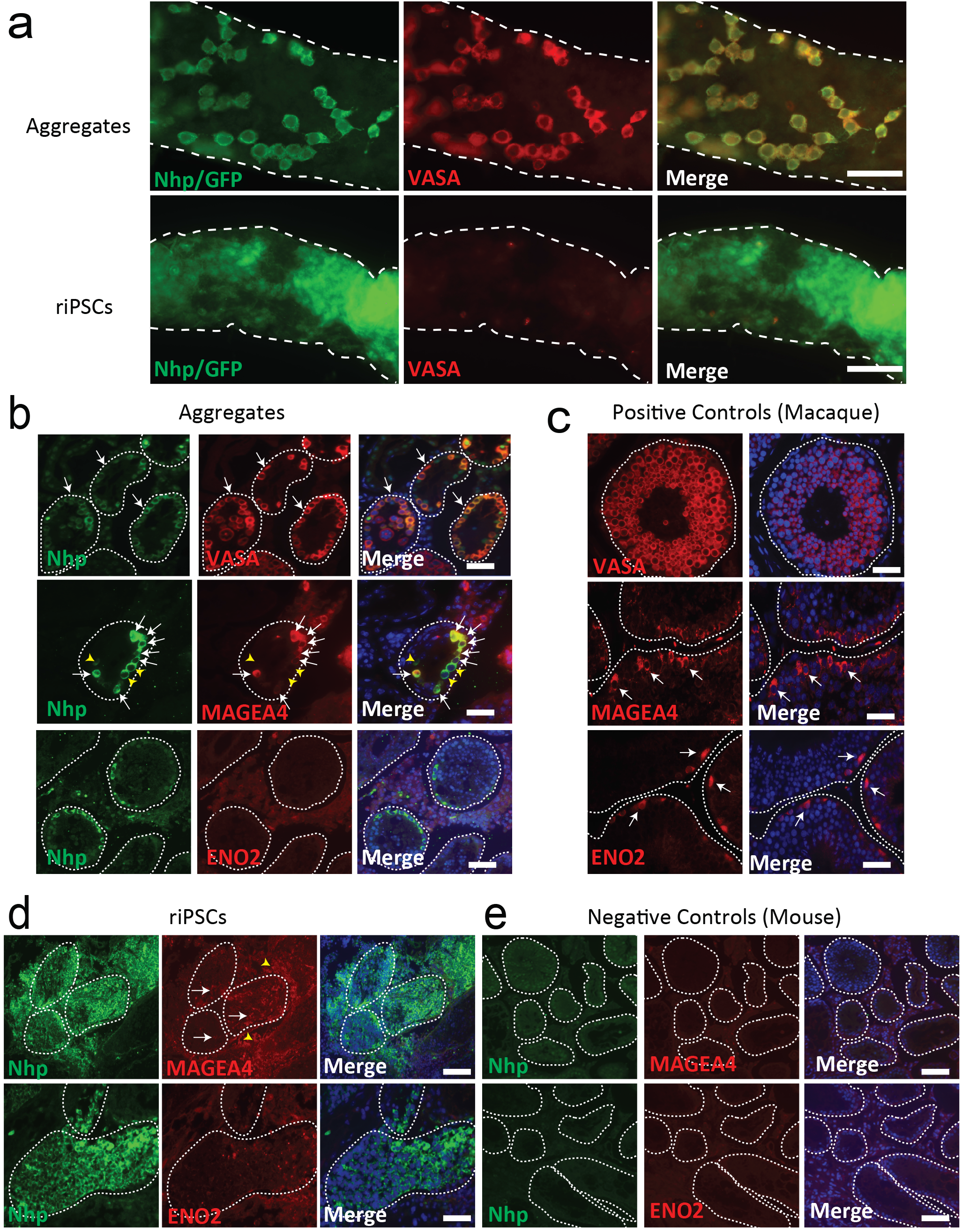
Xenotranplantation of rPGCLCs leads to the expression of the late stage PGC marker, VASA. (a) Whole mount IF staining of mouse testis seminiferous tubules that received day 8 (D8) aggregates containing PGCLCs, reveals the presence of NHP/GFP (green) cells, near the basement membrane of the tubules (white dotted outline) that coexpress the late-stage PGC marker VASA (red). While riPSCs recipient testis contained NHP/GFP positive cells that were VASA negative. Scale bars, 40μm. (b) IF staining on aggregate recipient, paraffin embedded, testis sections for NHP positive cells (green) reveals that these cells are found within the seminiferous tubules (white dotted lines) and express the germ cell marker VASA (white arrows). Also, a subset of NHP+cells express MAGEA4 (red, white arrows), while ENO2 is absent in NHP+ cells. Scale bars, 40μm. (c) Positive staining of the germ cell markers VASA (red), MAGEA4 (red) and ENO2 (red) in adult rhesus macaque testis. Scale bars, 40μm. (d) NHP positive cells coexpress MAGEA4 inside (white arrows) and outside (yellow arrowheads) the seminiferous tubules (white dotted lines). Scale bars, 40μm. (e) IF on control mouse testis reveal that they are negative for the primate marker NHP and the germ cell markers MAGEA4 and ENO2. Scale bars, 40μm.

To confirm that xenotransplantation supported hPGCLC differentiation to more advanced stages, sections were stained for VASA (the late-stage PGC marker) as well as the spermatogonial stem cell markers MAGEA4 and ENO2. Using these markers, our results demonstrate that almost all Nhp positive rPGCLCs in the recipient mouse testicular tubules differentiated into VASA positive germ cells confirming the results of the whole mount. Furthermore, these NHP-positive rPGCLCs were exclusively located on the basement membrane, typical of spermatogonia. A fraction (28%) of NHP positive rPGCLCs also expressed the spermatogonial gene MAGEA4 (Fig. 4b). In contrast, ENO2 was not expressed in any NHP-positive cells in the mouse testis (Fig. 4b) tubules. Given that only one of the two spermatogonial stem cell markers (MAGEA4) was expressed in the NHP-positive germ cells in the xenotransplants, we examined MAGEA4 (and ENO2) expression in embryonic rPGCs with the hypothesis that MAGEA4 may be expressed earlier in germ cell development prior to spermatogonia formation as was reported for humans^24,25^. Our data shows that MAGEA4 (but not ENO2) is expressed in rPGCs at the time when rPGCs are migrating through the dorsal mesentery, as well as, when making contact with the genital ridge epithelium (Supplementary Fig. 4f-g). Therefore the VASA+/MAGEA4+ rPGCLCs in the xenotransplants are most likely late-stage rPGCLCs and not spermatogonia given the lack of ENO2 expression. We also stained recipient testes transplanted with riPSCs and found large clumps of NHP-positive cells within the tubules that did not express MAGEA4 or ENO2 (Fig. 4d). However, we noted NHP-positive cells outside of the tubules in the interstitial area were MAGEA4 positive (Fig. 4d, yellow arrow heads).

Due to the inability of primate cells to undergo complete spermatogenesis in seminiferous tubules of mice, we also performed homologous transplantation of rPGCLCs into irradiated rhesus macaque male testicles to ascertain whether xenotransplantation was the barrier to rPGCLC differentiation into ENO2-positive spermatogonia. To achieve this, we transplanted dissociated aggregates containing rPGCLCs or undifferentiated riPSCs into the left and right testis, respectively, of rhesus macaque males depleted of germ cells by irradiation (n=2). The monkeys were also administered GnRH-antagonist treatment starting immediately after irradiation for 2 months until the time of transplantation. GnRH-antagonist treatment is expected to facilitate survival and colonization of transplanted cells^15^. Serum testosterone levels were monitored to confirm transient suppression of testosterone during GnRH antagonist treatment and maintenance of Leydig cell functions after irradiation (Fig. 5a). Ultrasound-guided rete testis injection was performed under general anesthesia to transplant 4.2 million single aggregate cells, and 9.8 million undifferentiated riPSCs into the rete testis. Following transplantation, testicular volumes and serum testosterone were measured monthly in each animal (Fig. 5a). In animal 9-174, the final volume and weight of recipient testicles at 7 months after transplantation were indistinguishable from each other, despite one testicle receiving aggregate cells, whereas the other testicle received undifferentiated riPSCs. In contrast, in the other animal (animal number 9-052), the testicle transplanted with undifferentiated riPSCs was 11-fold heavier relative to the contra-lateral testicle transplanted with aggregate cells (Fig. 5a).

**Fig. 5.**
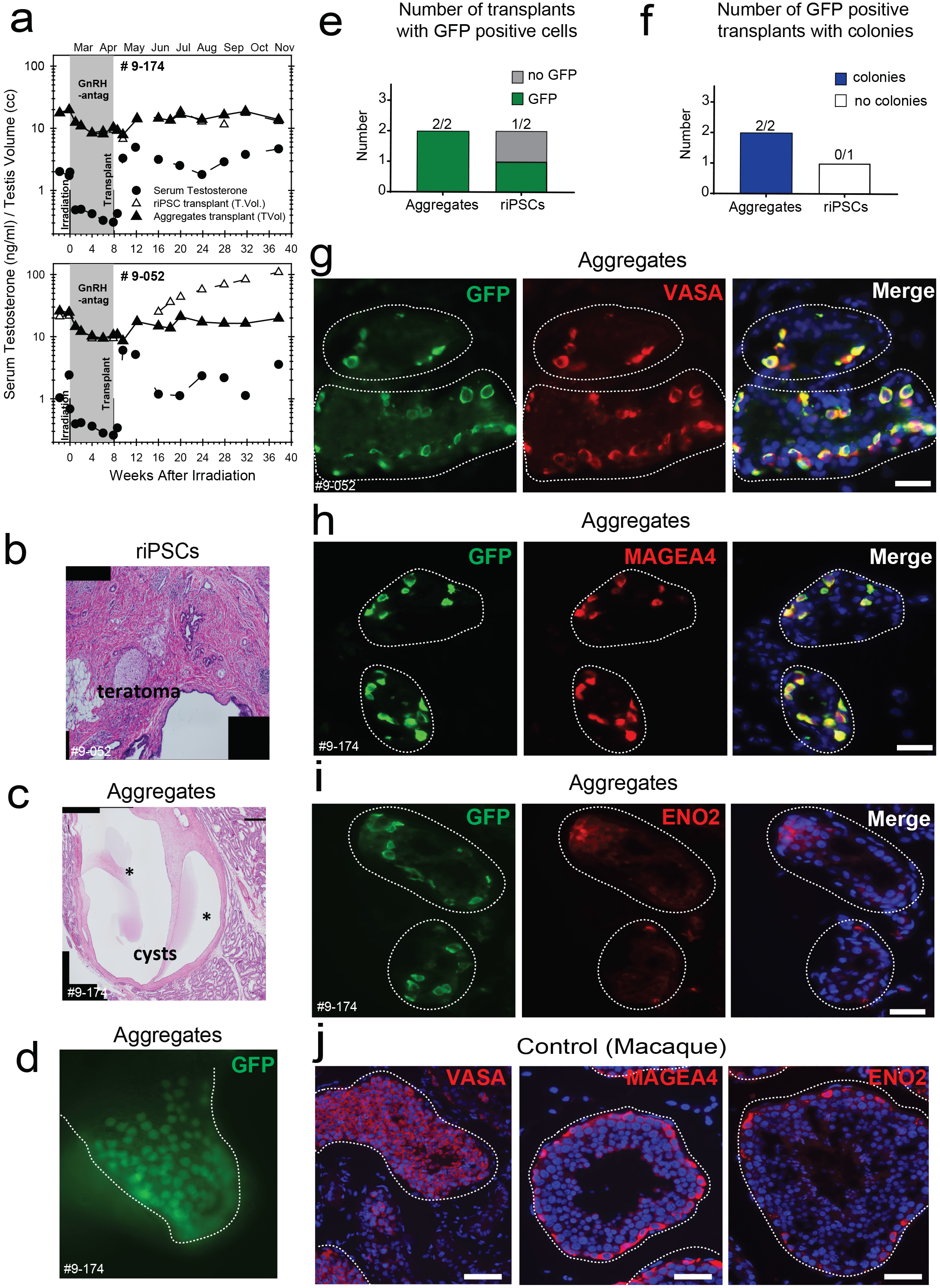
Homologous transplantation of rPGCLCs resulted in the expression of late stage PGC markers. (a) The volume of recipient testis that received aggregate donor cells returned to baseline levels, comparable to the volume prior to irradiation and subsequent GnRH antagonist treatment. Of the two testes that received iPSCs, one testis had increased volume, whereas the other testis did not. (b) H&E staining of riPSCs recipient testis reveals the formation of teratomas. (c) H&E staining of aggregate recipient testis reveals large cysts (asterisks) that form near normal tubules. (d) Expression of GFP in testis that received aggregate cells within the seminiferous tubules (white dotted lines). (e) A graph of the number of transplants that had GFP positive donor cells. (f) Graph of the number of GFP positive transplants with colonies. (g-j) IF on sections of rhesus recipient testis that received aggregate cells for donor cells (GFP positive, green) reveals that donor cells co-express the late stage PGC markers (g) VASA (red) and (h) MAGEA4 (red), but not the spermatogonial maker (i) ENO2. Nuclear staining using DAPI (blue) is shown in merged panels for g-i. Scale bars, 40μm. (j) Positive IF staining of rhesus adult testis for VASA (red), MAGEA4 (red), or ENO2(red) in combination with the nuclear marker DAPI (blue). Scale bars, 40μm.

Given the abnormal growth of the testicle in animal 9-052 that received undifferentiated iPSCs, the experiment was terminated at seven months after transplant, and the testicles from both animals were removed. Upon dissecting the testicles from animal 9-052, it was apparent under visual inspection that the testicle receiving aggregate cells contained cysts, whereas the testicle that received undifferentiated riPSCs was filled with solid tumorigenic masses (Supplementary Fig. 5a, 5b). It was also possible to identify GFP positive signal in both testicles from 9-052 (Supplementary Fig. 5c). Histological analysis confirmed that transplantation of undifferentiated riPSCs resulted in formation of a teratoma, which had almost completely invaded the testicle (Fig. 5b). The contralateral testicle that received aggregate cells, however, developed large epithelial cysts, with no evidence of teratomas (Fig. 5c, asterisks).

In order to examine whether rPGCLCs were capable of differentiation in the seminiferous tubule epithelium, the testicles from 9-174 and 9-052 were fixed, embedded and serially sectioned. Using GFP to mark the transplanted cells, we identified rare GFP positive cells along the seminiferous tubule basement membrane of rhesus macaque testicles injected with aggregates (2/2 transplants of aggregates) (Fig. 5d-e) that resulted in the formation of colonies (Fig. 5f). In contrast, the GFP-positive testicle transplanted with undifferentiated riPSCs exhibited no colonies (Fig. 5f). The GFP positive cells did not represent complete spermatogenesis, and instead were organized as individual cells, and occasionally pairs. Moreover, we show that the GFP positive cells along the basement membrane were positive for VASA and MAGEA4 but were negative for ENO2 consistent with the results obtained from the testicular xenotransplants (Fig. 5g-i). Control rhesus macaque seminiferous tubules confirmed antibody specificity for VASA, MAGEA4, and ENO2 (Fig. 5j). The GFP-positive testicle transplanted with undifferentiated riPSCs exhibited no evidence of VASA, MAGEA4, or ENO2 staining (data not shown). Taken together, these findings confirm homologous transplantation of aggregates containing rPGCLCs leads to differentiation into VASA positive, MAGEA4 positive cells, and that xenotransplantation is a useful indicator to predict the outcome of homologous transplantation into adult rhesus macaque testicles.

## Discussion

Here we establish the rhesus macaque as another non-human primate species that is amendable to rPGCLC differentiation *in vitro*. Consistent with previous studies in the human^1,2^ and in the cynomolgus macaque^4^, *VASA* RNA and protein are also not detected in PGCLCs during the aggregate stage of differentiation. VASA is an evolutionary conserved gene that is expressed in germline cells of all metazoan, and in mammals is expressed as PGCs approach the developing genital ridge^26^. In the current study, we show that almost all rPGCs at D28 are VASA positive, and located in the dorsal mesentery and the genital ridge epithelium, consistent with previous reports describing VASA protein expression in extra-gonadal human PGCs and early stage cynoPGCs^5,27^. We propose that the expression of VASA protein in rPGCLCs following either testicular xenotransplantation into the mouse or homologous testicular transplantation into the rhesus macaque is consistent with the *in vitro* model generating early rPGCLCs equivalent to *bona fide* PGCs in the embryo that are competent to differentiate into the VASA positive stage.

The mechanism responsible for inducing VASA protein in mammalian PGCs as they approach the genital ridge is unclear. Using transgenic mouse technology it was reported that expression of mouse vasa homologue (*mvh*) in mouse embryonic somatic cells was associated with loss of DNA methylation^28^. In the current study we show that DNA methylation levels are high in rPGCLCs during the aggregate stages, which is consistent with the hypothesis that VASA expression could be regulated by DNA methylation.

Loss of DNA methylation in cynoPGCs occurs before VASA expression at day 21 of embryo development with PGC specification occurring at day 11. Therefore, our data suggests that the current approach for differentiating rPGCLCs in aggregates supports the formation of newly specified rPGCLCs but does not support the differentiation into the globally demethylated or VASA positive state. Given that rPGCLCs numbers are lost between day 8 and 15 of aggregate differentiation *in vitro*, we propose that it is the transfer of rPGCLCs to a new environment supportive of rPGCLC survival that is primarily responsible for advancing rPGCLC differentiation into VASA-positive PGCs. Consistent with this hypothesis, culture of mPGCs or mPGCLCs in growth factors and chemicals that maintain PGC survival also enable the transition from MVH-negative to MVH-positive states^29,30^. Therefore, it is likely that the adult testicular cells *per se* are not responsible for inducing VASA expression in the rPGCLCs, but instead require a supportive environment that enables the rPGCLC survival. Given this result, we anticipate that chemically defined conditions could be identified in the future to support the differentiation of rPGCLCs into VASA positive germ cells *in vitro* without the requirement for transplantation.

In addition to VASA, we discovered that the spermatogonial marker MAGEA4 was expressed in germ cell colonies arising from the transplanted aggregates cells.

MAGEA4 is not only a spermatogonial stem cell marker, it is also expressed in human germ cell tumor cells such as in cells of classical seminomas. Here we show that MAGEA4 is expressed much earlier, being present in rPGCs by D28, as they approach the genital ridge epithelium. Given that we do not know when ENO2 is expressed during the later gestational stages, or whether it is exclusive to adult spermatogonia, it is difficult to predict whether the lack of ENO2 in the rPGCLCs is because of a block in developmental timing from a MAGEA4 to an ENO2 positive spermatogonia, or whether *ENO2* is abnormally silenced in the rPGCLCs. Our RNA-Seq data at Carnegie Stage 23 suggest that *ENO2* RNA is expressed at low to background levels in rPGCs. Future studies aimed at characterizing the pro-spermatogonial stage of primate PGC differentiation will be critical to understanding the lack of ENO2 expression following transplantation.

Previous studies have shown that testicular xenotransplantation of undifferentiated hESCs and hiPSCs into the testicles of busulfan-treated nude mice result in the differentiation of both VASA positive PGCs accompanied by teratoma, embryonal carcinoma and yolk sac tumor formation^31^. The tumors that arose in these studies most likely originate from hiPSCs/hESCs because testicular xenotransplantation of embryonic testes containing bona fide PGCs results in colony formation without tumorigenesis^16^. Unlike studies using human pluripotent stem cells, we did not identify any evidence of VASA positive cells in the testes transplanted with undifferentiated riPSCs. However, MAGEA4-positive, VASA-negative cells were observed. MAGEA4 is a member of the cancer/testis antigen group, and has been proposed as a novel somatic cell cancer therapy target^32,33^. Given the widespread expression of MAGEA4 in cancer cells, we hypothesize that the MAGEA4+/VASA-negative cells originating from the transplanted riPSCs correspond to MAGEA4+ tumor cells. Although xenotransplantation of aggregates containing rPGCLCs did not result in teratomas, these transplants yielded cysts in recipient testes. In future studies purified PGCLCs could be transplanted to determine whether the cyst cells are originating from the PGCLCs, aggregate somatic cells, or both. Notably, these cysts were also observed following homologous aggregate cell transplantation into the rhesus macaque testis indicating that this outcome is not simply due to species specific differences between transplanted cells and recipient.

Successful resumption of spermatogenesis from donor cells in homologous monkey-to-monkey transplants involves transfer of tissues containing adult spermatogonia^17,19^. In the current study, the lack of spermatogenesis in homologous transplants could be due to either a poor quality rPGCLC or alternatively the possibility that rPGCs are incapable of differentiating in an adult niche. In mouse transplantation studies where mPGC transplantation was successful, neonatal infertile recipient mice were capable of supporting spermatogenesis from PGCs, whereas adult infertile recipients support spermatogenesis only when transplanted with tissues containing pro-spermatogonia or spermatogonia^8^. The only way to determine whether rPGCs have the competency to fully differentiate into haploid sperm would be to perform homologous transplants of bona fide rPGCs into infertile rhesus recipients at different ages. The feasibility of this theoretical experiment is challenged by the very low number of PGCs that are found during the embryonic stages of interest.

In summary, we show that rPGCLCs differentiated *in vitro* are competent to differentiate into VASA positive germ cells. Our studies also suggest that testicular xenotransplantation may be a cost-effective option for testing lineage commitment of *in vitro* specified germs cells prior to conducting more costly, and time consuming nonhuman primate, autologous or homologous transplantation experiments. Although it i s well known that full reconstitution of spermatogenesis cannot be achieved in the mouse from non-human primate cells, our study highlights the similarities when either mice or monkeys are used as recipients of *in vitro* differentiated PGCLCs.

## Methods

### Time-Mated Breeding of Rhesus Macaque

Time-mated breeding of rhesus macaque males and females was performed by measuring estradiol daily in the female starting from day 5 to 8 after menses began as previously described^16^. Pregnancy was confirmed by measuring progesterone as well as by ultrasound. D28 embryos were collected by C-section. All rhesus macaque time-mated breeding experiments were conducted following approval of the ONPRC Institutional Animal Care and Use Committee.

### Maintenance and Directed Differentiation of riPSC89 and riPSC90

Undifferentiated riPSC89 and riPSC90 cells were cultured on a feeder layer of mitomycin C-treated MEFs in human Embryonic Stem Cell media (DMEM/F-12 (Life Technologies), 20% KSR (Life Technologies), 10 ng/mL bFGF (R & D Systems), 1% nonessential amino acids (Life Technologies), 2 mM L-glutamine (Life Technologies), Primocin™ (Invivogen), and 0.1 mM β-mercaptoethanol (Sigma)). Media was changed daily and colonies were passaged manually every 5 days. For iMeLC inductions we followed previously published protocols with minor modifications^2^. Briefly, Day 5 riPSCs colonies were trypsinized (0.05% Trypsin (LifeTechnologies) and resuspended in MEF media. The MEFs were depleted by plating the cell suspension in tissue culture dishes, two times, for 5 minutes each. The resulting cell suspension was pelleted and resuspended in iMeLC media (GMEM (Life Technologies), 15% KSR (Life Technologies), 0.1 mM nonessential amino acids (Life Technologies), Penicilin/Streptomycin/L-Glutamine (Life Technologies), Primocin™ (Invivogen), 0.1 mM β-mercaptoethanol (Sigma), sodium pyruvate(Life Technologies), activin A (PeproTech), CHIR99021(Stemgent), Y-27632 (Stemgent)), filtered through a 40 μm cell strainer (Falcon) and plated at a density of 1.0 x 10^5^ cells per well of a human plasma fibronectin (Invitrogen)-coated 12-well plate.

After 24 hours of incubation at 37°C with 5.0% CO2, iMeLCs were trypsinized (0.05% trypsin, Life Technologies) and resuspended in PGCLC media (GMEM (Life Technologies), 15% KSR (Life Technologies), 0.1 mM nonessential amino acids (Life Technologies), Penicilin/Streptomycin/L-Glutamine (Life Technologies), Primocin™ (Invivogen), 0.1 mM β-mercaptoethanol (Sigma), Sodium pyruvate (Life Technologies), 10 ng/mL human LIF(EMD Millipore), 200 ng/mL BMP4 (R&D systems), 50 ng/mL EGF (Fisher Scientific), 10 μM Y-27632 (Stemgent)), and plated at a density of 3.0 x 10^3^ cells per well of a low adherence spheroid forming 96-well plate (Corning). Aggregates were collected for analysis on Day 1, 2, 3, 4, 8 and 15 of directed PGCLC differentiation.

### Immunofluorescence staining

Aggregates containing PGCLCs were collected then fixed in 4% PFA and embedded in histogel (ThermoScientific) to facilitate subsequent embedding into paraffin blocks. Sections of aggregates (5 μm) placed onto microscope slides were then de-paraffinized and rehydrated through a xylene, ethanol series. For antigen retrieval, slides were heated to 95°C in Tris-EDTA solution (10mM Tris Base, 1mM EDTA Solution, 0.05% Tween-20, pH 9.0). Sections were permeabilized (PBS, 0.05% Triton X-100) and then blocked in PBS containing 10% normal donkey serum. Primary antibodies (Table 1) were incubated overnight at 4°C, followed by incubating with secondary antibodies at room temperature (Table 1). Mounting media (Prolonggold anti-fade w/DAPI, Invitrogen) was added and slides were sealed. Slides were allowed to cure for at least 24 hours at 4°C prior to imaging.

**Table 1.**
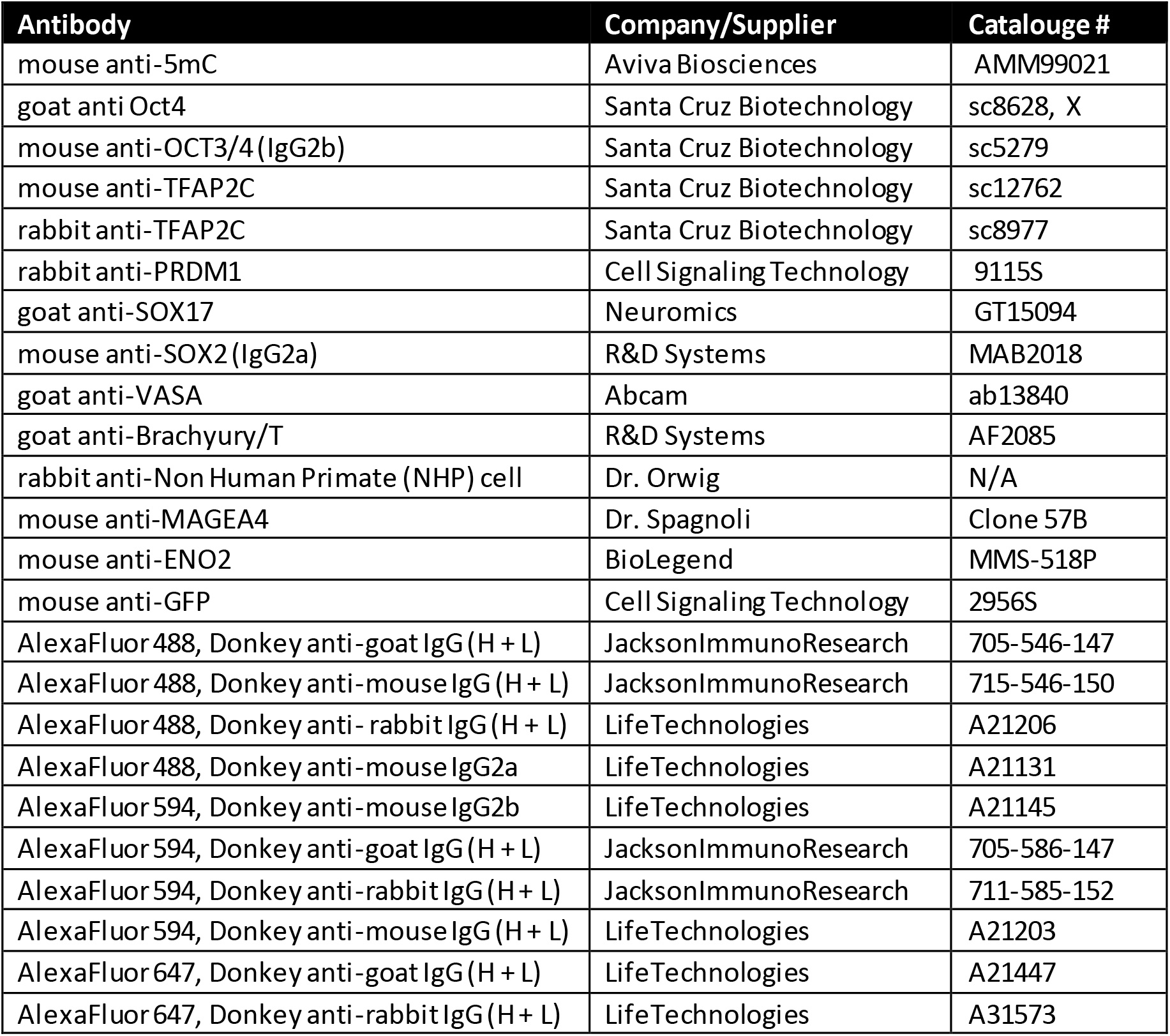
List of antibodies used for experiments

riPSCs and riMeLCs were plated onto culture slides (Corning) and grown overnight in their respective media, at 37C with 5.0% CO2. Cells were washed with PBS then fixed in 4% PFA for 10 minutes. Cells were rinsed with PBST and then permeablized (PBS, 0.05% Triton X-100) for 10 minutes. Nonspecific binding was blocked (PBS, 10% normal Donkey Serum for 30 minutes) and then incubated overnight at 4°C with primary antibodies (Table 1). Species-specific secondary antibodies were added to slides and incubated at room temperature for 1 hour. Mounting media was added, slides were sealed, and then cured for 24 hours.

Mouse testes tissues were fixed overnight in 4% paraformaldehyde, 5μm thick paraffin sections were used for the immunofluorescence study. The monkey testes were fixed in cold 4% paraformaldehyde overnight, processed through a sucrose gradient and then embedded in OCT; 8 μm cryosections were cut for immunofluorescence staining. For testicular histology of both mouse and monkey, Bouin’s or 4% paraformaldehyde-fixed tissues were embedded in paraffin and sections were stained in periodic acid-Shiffs-hematoxylin.

For fluorescent immunostaining, either the deparaffinized or frozen sections were subjected to antigen retrieval by initially heating in boiling citrate buffer (BioGnenex) for 2 min followed by cooling for 30-60 min. The slides were rinsed and blocked in antibody diluent. Subsequently, sections were stained with the following primary antibodies in antibody diluent: goat anti-VASA, mouse anti-ENO2, mouse anti-MAGE-A4 (Clone 57B, kindly provided by Dr. Giulio Spagnoli, University Hospital Basel, Switzerland), rabbit anti-GFP, rabbit anti-rhesus testis cell (NHP) (provided by Kyle Orwig). For double immunofluorescent staining, the two primary antibodies incubated with the tissue sections were detected with the species-specific secondary antibodies (Table 1).

Stained sections were mounted with VectaShield mounting media containing DAPI (Vector Laboratories) and imaged. Positive immunoreactivity was validated by omission of primary antibody and testing the antibody in a tissue in which it is known to be positive.

### Microscopy

Confocal images of the riPSC89 line, riPSC90 line, iMeLCs, and sectioned aggregates were examined on an LSM 880 (Carl Zeiss) with a Plan-Apochromat 20×/0.8NA and a Plan-Apochromat 63×/1.4NA M27 oil immersion objective at room temperature. Acquired images were processed using IMARIS 8.1 (Bitplane, Zurich, Switzerland). H&E slides were examined on an Olympus BX-61 light microscope. Images were processed with Image J version 1.51d (NIH). D28 embryos, teratomas, and cysts images were stitched together using the grid-stitching plugin in Image J version 1.51d (NIH).

For immunofluorescence imaging of tissue sections or whole mounts of seminiferous tubules, a Leica DM 4000B (Leica Microsystems) microscope was used.

### Fluorescence activated cell sorting analysis of rPGCLCs

Day 4 and Day 8 PGCLCs were dissociated using 0.05% trypsin or 0.1 mg/mL Collagenase,Type IV (Gibco)/ 0.25% trypsin, respectively, and re-suspended in FACS buffer. Dissociated cells were incubated with anti-ITGA6-BV421 (Fisher Scientific) and EpCAM-PE (LifeTechnologies) antibodies. Double positive cells were collected using an ARIA-H Fluorescence Activated Cell sorter. Cells were sorted into RLT buffer and stored at −20°C until ready to isolate RNA. Sorts of rPGCLCs were performed on at least 3 replicates for each group.

### Establishment of the riPSC89^UbiC:GFP^ reporter line

The riPSC89 GFP-reporter line was established through episomal delivery of the lentivirus “GFP-IRES_PURO_cassette”, a kind gift from Dr. Zoran, using lipofectamine 3000 (Invitrogen). Karyotypes performed by CellLineGenetics (Madison, WI) indicated that the GFP-expressing riPSC89 line had a normal karyotype.

### Whole mount Immunofluorescence staining of seminiferous tubules

For quantitative analysis of donor rhesus testis cell colonization, intact seminiferous tubules were prepared from nude mouse recipient testes, collected 8 weeks after transplantation as previously described^15^. Donor-derived colonies of spermatogonia were detected in intact seminiferous tubules by whole-mount immunofluorescent staining with the rhesus testis cell antiserum^17^. Samples were dehydrated stepwise in methanol and then incubated in MeOH:DMSO:H2O2 (4:1:1) for 2 hours. The rhesus testis-cell antibody was used and detected with rabbit secondary antibodies (Table 1). Samples were mounted with Vectashield medium containing DAPI (Vector Laboratories) on slides with raised coverslips and visualized by fluorescence microscopy.

In co-staining experiments, the rhesus testis cell and VASA antibodies were detected with species-specific antibodies (Table 1).

### RT-PCR

Purified RNA was extracted from undifferentiated riPSCs, riMeLCs, day 4 and day 8 PGCLC sorted cells(EpCAM^+^/ITAG6^+^) using the RNAeasy mini kit (Qiagen). RNA was converted to cDNA using Superscript II Reverse Transcriptase (Invitrogen). Gene expression analysis was performed using the following gene expression assays: TFAP2C: rh02844868_m1 (Applied Biosystems); PRDM1: rh02837836_m1(Applied Biosystems); KIT: qMccCIP0023334(BioRad); NANOS3: qMccCEP0031201(BioRad); GAPDH:qMccCIP0038690(BioRad).

### Xenotransplantation of rPGCLCs and riPSCs into the seminiferous tubules

Day 8 rPGCLCs aggregates (0.1 mg/mL Collagenase,Type IV (Gibco) then 0.25% trypsin (Gibco) and riPSCs (0.1 mg/mL Collagenase, Type IV (Gibco) were dissociated prior to transplantation. They were injected in the right and left testis of nude mice depleted of germ cells by irradiation using a^137^Cs gamma-ray unit. The radiation was delivered as an initial 1.5-Gy dose and followed by a second dose of 12 Gy^34,35^. As previously described^12^, about 7μL of solution (a range of cells from 5.4 X 10^4^ to 2.7 x 10^5^ cells) was injected per testis. Mice were allowed to recover and sacrificed 2 months later. In most cases, testes were fixed in 4% paraformaldehyde, paraffin-embedded and sectioned for immunofluorescence analysis. In some cases testis seminiferous tubules were dispersed using collagenase and DNase, fixed in 4% paraformaldehyde and prepared for whole mount immunofluorescence staining.

### Homologous transplantation rPGCLCs and riPSCs into the monkey seminiferous tubules

Day 8 rPGCLCs aggregates (0.1 mg/mL Collagenase,Type IV (Gibco) then 0. 25% trypsin (Gibco)) and riPSCs (0.1 mg/mL Collagenase, Type IV (Gibco) were dissociated prior to transplantation. Both rPGCLCs and riPSCs were suspended in MEMα containing 10% FBS, trypan blue (0.4 mg/mL; Sigma-Aldrich), 20% (v/v) Optison ultrasound contrast agent (GE Healthcare, Waukesha,WI, USA), 1% antibiotic-antimycotic (a combination of penicillin, streptomycin and amphotericin B; Gibco) and DNase I (0.1 mg/mL) in a total volume of 0.5 mL. Using ultrasound guidance, to locate the rete testis, riPSCs were injected into the right testis and rPGCLC to the left testis of Rhesus macaques depleted of germ cells by testicular irradiation with 7 Gy^15^. Either 9.8 million riPSCs or 4.2 million rPGCLCs were injected per testis. To possibly enhance the grafting of the transplanted cells the monkeys were given hormone suppression treatment using a GnRH-antagonist Acyline (obtained from the Contraceptive Development Program of the NICHD, Rockville, MD, USA) for 2 months starting immediately after irradiation and until the time of transplantation^15^. To prevent T cell-mediated rejection of the grafted cells, transplant recipients were treated with human/mouse chimeric anti-CD154 IgG 5C8 (NIH Nonhuman Primate Reagent Resource, University of Massachusetts Medical School, Boston, MA) at 20 mg/kg on days -1, 0, 3, 10, 18, 28, and monthly thereafter. Changes in testes sizes and serum testosterone levels were monitored during the post-transplantation period and the testes were harvested 7 months after transplantation. The sliced testes were fixed in 4% paraformaldehyde and after initial observation under fluorescence scope for any GFP signal, were either embedded in OCT and cryosectioned for immunofluorescence analysis, or embedded in paraffin for histology.

## Acknowledgements

EJR was supported by the National Institute of General Medical Sciences of the National Institutes of Health under award number R25GM055052 awarded to T. Hasson. Rhesus embryo collections were supported by the grant P51 OD011092 (JDH and ATC). Funding for riPSCs line establishment, cell culture, and transplantation experiments were funded with the support of the grant P01HD075795 (GS, MLM, KEO, ATC).

## Author Contributions

E.S. conceived the experiments, performed experiments and wrote the manuscript; D.C. performed FACS and RNA-seq experiments on rPGCs and rPGCLCs; E.J.R. performed Immunofluorescence experiments on rPGCs and rPGCLCs; Z.W. and T.N.L performed xenotransplantation experiments; K.A.P. and J.M.M. helped with homologous transplantations in rhesus macaques; R.C.T performed dose and radiation field planning on each individual monkey; J.D.H. conceived experiments maintained rhesus macaque IACUC approval for time mated breeding’s, and oversight of rhesus macaque work in Oregon; M.L.M performed experiments, maintained IACUC approval, and oversight of mouse xenotransplantation and rhesus macaque transplantation in Houston; K.E.O. conceived the experiments, performed homologous rhesus macaque transplants; G.S. conceived and performed experiments, maintained IACUC approval, and oversight of mouse and rhesus macaque work in Houston; A.T.C. conceived the experiments, performed experiments, maintained Institutional Biosafety Approval for rhesus macaque riPSCs/rPGCLCs work and wrote the manuscript.

**Additional information**

**Supplementary Information**

**Competing financial interests**

